# SpatialDE - Identification of spatially variable genes

**DOI:** 10.1101/143321

**Authors:** Valentine Svensson, Sarah A Teichmann, Oliver Stegle

## Abstract

Technological advances have enabled low-input RNA-sequencing, paving the way for assaying transcriptome variation in spatial contexts, including in tissues. While the generation of spatially resolved transcriptome maps is increasingly feasible, computational methods for analysing the resulting data are not established. Existing analysis strategies either ignore the spatial component of gene expression variation, or require discretization of the cells into coarse grained groups.

To address this, we have developed SpatialDE, a computational framework for identifying and characterizing spatially variable genes. Our method generalizes variable gene selection, as used in population-and single-cell studies, to spatial expression profiles. To illustrate the broad utility of our approach, we apply SpatialDE to spatial transcriptomics data, and to data from single cell methods based on multiplexed *in situ* hybridisation (SeqFISH and MERFISH). SpatialDE enables the statistically robust identification of spatially variable genes, thereby identifying genes with known disease implications, several of which are missed by conventional variable gene selection. Additionally, to enable gene-expressed based histology, SpatialDE implements a spatial gene clustering model which we call “automatic expression histology,” allowing to classify genes into groups with distinct spatial patterns.

---

Technological advances have helped to miniaturize and parallelize genomics, thereby enabling high-throughput transcriptome profiling from low quantities of starting material, including in single cells. Increased experimental throughput has also fostered new experimental designs, where the spatial context of gene expression variation can now be directly assayed, which is critical for characterizing complex tissue architectures in multicellular organisms. The spatial context of gene expression is crucial for determining functions and phenotypes of cells^1,2^. Spatial expression variation can reflect communication between adjacent cells, or can be caused by cells that migrate to specific locations in a tissue to perform their functions.

Several experimental methods to measure gene expression levels in a spatial context have been established, which differ in resolution, accuracy and throughput. These include the computational integration of single cell RNAseq data with a spatial reference dataset^3,4^, careful collection and recording of spatial location of samples^5^, parallel profiling of mRNA using barcodes on a grid of known spatial locations^5–7^, and methods based on multiplexed *in situ* hybridization^8,9^ or sequencing^10–12^.

A first critical step in the analysis of the resulting datasets is to identify the genes that exhibit spatial variation across the tissue. However, existing approaches for identifying highly variable genes^13,14^, as in single-cell RNA-sequencing (scRNA-seq) studies, ignore the spatial location and hence do not measure *spatial* variability (**Figure 1A**). Alternatively, researchers have applied ANOVA to test for differential expression between groups of cells, either derived using *a priori* defined (discrete) cell annotations, or based on cell clustering^3,4,7,8,10^, with some methods incorporating spatial information^15^. Importantly, such strategies fall short of detecting variation that is not well captured by discrete groups, including linear and nonlinear trends, periodic expression patterns and other complex patterns of expression variation.

**Figure 1.**
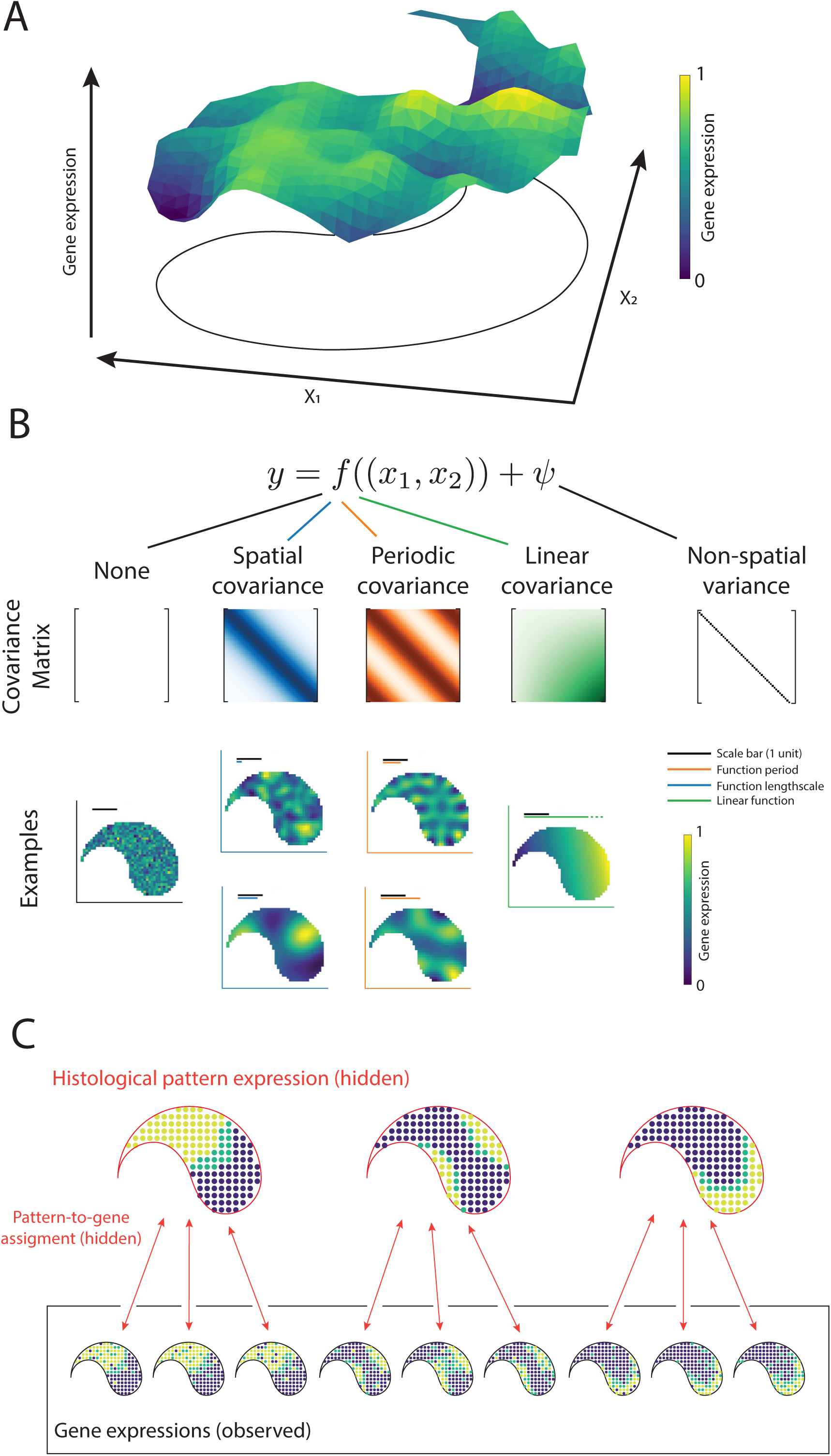
Overview of SpatialDE for the identification of spatially variable genes. (**A**) In spatial gene expression studies, expression levels vary in ways that depend on spatial coordinates. SpatialDE defines spatial dependence for a given gene using a non-parametric regression model, testing whether gene expression levels at different locations co-vary in a manner that depends on their relative location. (**B**) SpatialDE partitions the expression variation into a spatial component (using functional dependencies f(x,y)), characterized by alternative spatial covariances, and observation noise (Psi). Alternative spatial covariance models considered by SpatialDE: no spatial effect (null model), general spatial, periodic spatial patterns, and linear trends. Representative simulated expression patterns are plotted below the corresponding covariance matrices. (**C**) Automatic expression histology models the expression levels of spatially variable genes using a set of unobserved tissue structure patterns. Both the underlying patterns and the gene-pattern assignments are learned from data.

To address this, we here propose a computational approach termed *SpatialDE* for identifying and characterizing *spatially variable* genes (SV genes). Our method builds on Gaussian process regression, a class of models that is widely used in geostatistics, also known as Kriging^16^. Briefly, for each gene, SpatialDE decomposes the expression variability into a spatial and non-spatial component (**Figure 1A-B**). These variance components are modelled using two random effect terms: a spatial variance component term that describes gene expression covariance as a function of the pairwise distance of cells and a second (noise) term that accounts for non-spatial expression variability. The ratio of the variance explained by these components quantifies the Fraction of Spatial Variance. Significant SV genes can be identified by comparing this full model to a model that assumes no spatial dependency of expression variation (**Figure 1B**, **Methods**).

In addition to identifying spatially variable genes, SpatialDE classifies the spatial patterns of individual genes thereby distinguishing between linear trends, periodic expression profiles or general spatial dependencies (**Figure 1B**). By interpreting the fitted model parameters it is possible to identify the length scale (the expected number of changes in direction in a unit interval^16^) or the period length of spatial patterns for individual genes (**Figure 1B, Supplementary Methods**). Finally, SpatialDE implements a spatial clustering method, *automatic expression histology* (AEH), within the same Gaussian process framework as used to test for SV genes, thereby identifying sets of genes that mark distinct spatial expression patterns (**Figure 1C**).

The SpatialDE is computationally efficient by leveraging computational shortcuts for efficient inference in linear mixed models^17^ and precomputing operations possible due to the structure of massively parallel molecular assays (**Methods**, **Supp. Fig. 1**). Taken together, SpatialDE is a widely applicable tool for the analysis of spatial transcriptomics datasets.

First, we applied our method to existing spatial transcriptomics data from mouse olfactory bulb^7^. Briefly, spatial transcriptomics gene expression levels were derived from thin tissue sections of frozen material, placed on an array with poly(dT) probes and spatially resolved DNA barcodes in a grid of circular “spots” with a diameter of 100 μm, thereby measuring mRNA variation with a resolution of 10-100 cells per spot (depending on tissue type). Following permeabilization, the mRNA is captured by the probes, and the spatial location can be recovered from sequenced barcodes. The resulting gene expression profiles can be analysed in combination with hematoxylin and eosin (HE) stained microscopic images of the tissue (**Figure 2A**).

**Figure 2.**
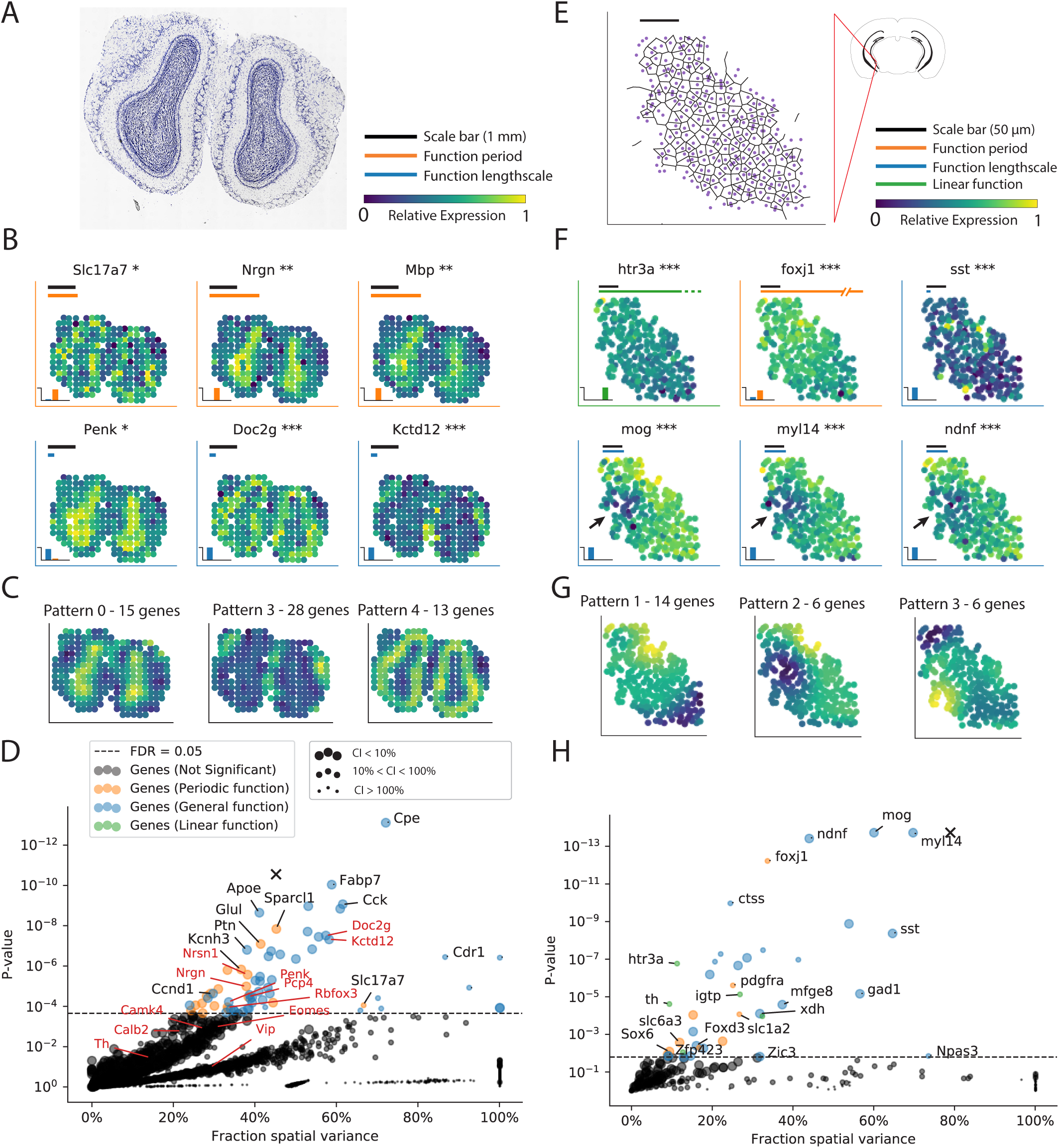
Applications of SpatialDE to spatial transcriptomics and data generated using SeqFISH. (**A**) Hematoxylin and eosin (HE) image for mouse olfactory bulb data from Stahl *et al.* (**B**) Visualization of six selected spatially variable genes (out of 67, FDR < 0.05). The black scale bar corresponds to 1 mm. For genes identified with periodic dependencies, the orange bar shows the fitted period length on the same scale. Analogously, the blue bar shows the fitted length scale for genes with general spatial trends. 2D plots show relative expression levels for genes across the tissue section coded in color. Stars next to gene names denote significance levels (* FDR < 0.05, ** FDR < 0.01, *** FDR < 0.001) of spatial variation. Insets in lower left show the posterior probability of the three functional classes for each gene (general spatial, periodic spatial, linear trend). (**C**) Three example histological expression patterns identified by automatic expression histology analysis, which correspond to visually distinct layers in the HE image. Color denotes the level of expression of the inferred patterns. The number of genes assigned to the respective patterns are noted in plot title. (**D**) Fraction of variance explained by spatial variation (FSV, x-axis) versus P-value (y-axis) for all genes. Dashed line corresponds to the FDR=0.05 significance level (N=67 genes). Genes classified as periodically variable are shown in orange (N=19), genes with a general spatial dependency in blue (N=48). Genes highlighted as classical histological markers in Stahl *et al* are displayed in red text, representative examples of SV genes are annotated with black text. Size of points indicates the uncertainty of FSV estimates with larger points corresponding to genes with smaller standard errors. The X symbol shows the result of applying SpatialDE to the estimated total RNA content per spot. (**E**) SeqFISH data from a region of mouse hippocampus from Shah *et al*^8^. Black scale bar corresponds to 50 pm, with Voronoi tessellation representative of tissue structure. (**F**) Expression patterns of six selected SV genes (out of 32, FDR < 0.05). Shown are genes with linear *(htr3a),* periodic (*foxj1*), and general spatial trends. Black arrows indicate distinct region of low expression of *Mog, Myl14* and *Ndnf.* (**G**) Three examples of histological expression patterns. (**H**) Proportion of variance (x-axis) versus P-value (y-axis) for 249 genes, as in D. Genes with a linear dependency are colored in green.

The SpatialDE test identified 67 SV genes (FDR < 0.05, **Supp. Table 1**) with clear spatial substructure, consistent with the matched HE stained image (**Figure 2A-B**). These included canonical marker genes highlighted in the primary analysis by Stahl *et al*^7^, such as *Penk, Doc2g,* and *Kctd12,* but also additional genes that define the granule cell layer of the bulb. Genes in the latter set were classified as periodically variable with period lengths corresponding to the distance between the centers of the hemispheres, including *Kcnh3*, *Nrgn,* or *Mbp* with 1.8 mm period length (**Figure 2B,** further examples in **Supp. Fig. 2**). Other genes with periodic patterns, such as the vesicular glutamate transporter *Slc17a7,* were identified with shorter periods (1.1 mm), and inspection revealed regularly dispersed regions, potentially identifying a pattern associated with higher neuron density^18^, suggesting that periodic expression patterns in tissues can be of considerable biological interest. Applying automatic expression histology in SpatialDE identified five canonical expression patterns, clearly demarcating structures visible in the HE image (**Figure 2C, Supp. Fig. 3A**).

As a second application, we considered tissue slices from breast cancer biopsies^7^, profiled using the same ST protocol (**Supp. Fig. 4**). SpatialDE identified 115 SV genes (FDR < 0.05), including seven genes with known roles in the disease that were highlighted in the primary analysis of the data (**Supp. Fig. 4B-C**). Significantly SV genes were enriched for collagens, which distinguish tissue substructure^19^ (Reactome term “Collagen formation,” P < 5 * 10^−14^ using gProfiler^20^, **Supp. Table 1**). Additionally, we identified the autophagy related gene *TP53INP2,* surrounding the structured tissue (**Supp. Fig. 4C**). Interestingly, the set of SV genes also included the cytokines *CXCL9* and *CXCL13,* both of which are expressed in a visually distinct region (**Supp. Fig. 4A**, black arrow), together with the IL12 receptor subunit gene *IL12RB1,* indicating a potential tumour related immune response in the tissue. Notably, neither of these genes (and N=29 others), were identified as differentially expressed when applying unsupervised clustering in conjunction with an ANOVA test between the identified groups of cells (**Supp. Fig. 5**). Furthermore, these genes do not have a high rank based on conventional highly variable genes measures (such as the mean-CV^2^ relation^13^ or mean-dropout relation^21^), which do not take the spatial context into account (**Supp. Fig. 6**).

Automatic expression histology of the SV genes in the breast cancer biopsy (**Supp. Fig. 3B**) most clearly separated the adipocytic from the denser region of the sample, but additionally identified a small region overlapping the tumour feature in the HE image. Among the 17 genes assigned to this pattern were the cytokines and cytokine receptors *CXCL9, CXCL13, IL12RB1,* and *IL21R* (**Supp. Table 1**).

Overall, variable genes detected by SpatialDE are complementary to existing methods. In particular, SpatialDE identifies genes with localized expression patterns, as indicated by small fitted length scales, or periodic patterns, which are not detected by methods that ignore spatial contexts (**Supp. Fig. 5E**). We confirmed the statistical calibration and the robustness of SpatialDE using randomized data (**Supp. Fig. 7**) and simulation experiments (**Supp. Fig. 8**).

Taken together, these results demonstrate that SpatialDE can be used to characterize biologically and potentially clinically relevant features in spatial tissue samples in the absence of *a priori* histological annotation.

SpatialDE is not limited to sequencing technologies, and can be applied to any expression data type with spatial and/or temporal resolution. To explore this, we applied the method to data generated using multiplexed single molecule FISH (smFISH), a method that allows to quantify gene expression with subcellular resolution for a large number of target genes in parallel. Briefly, probes are sequentially hybridized to RNA while carrying barcodes of fluorophores, which allows to quantify gene expression of up to several thousand probes^22^ using high-content imaging.

We applied SpatialDE to multiplexed smFISH data of cells from mouse hippocampus, generated using SeqFISH^8^. This study considered 249 genes that were chosen to investigate the cell type composition along dorsal and ventral axes of the hippocampus (**Figure 2E**). SpatialDE identified 32 SV genes (FDR < 0.05), with the three highest ranking genes: *Mog, Myl14,* and *Ndnf* displaying a distinct region of lower expression (**Figure 2F**, black arrows). Again, SpatialDE identified genes with different types of spatial variation, including linear trends (N=5) and periodic patterns (N=8, **Figure 2H**, additional examples in **Supp. Fig. 9**) and these genes could be grouped into histological expression patterns using the AEH method (**Figure 2G, Supp. Fig. 2C**). Visual inspection of all the 249 genes supports the ranking of spatial variation from SpatialDE (**Supp. Fig. 10**).

SpatialDE can also be used to test for spatial expression variation in cell culture systems, where spatial variation typically is not expected *a priori*. As an example, we considered data from another multiplexed smFISH dataset generated using MERFISH with 140 probes on a human osteosarcoma cell line^9^ (**Supp. Fig. 11A-B**). In the primary analysis surprisingly *Moffitt et al* discovered spatially restricted populations of cells with higher proliferation rates. Interestingly, our model detected that a substantial proportion of the genes assayed were spatially variable (N=92, 65%, FDR<0.05), which recapitulates the results from the primary analysis. Indeed, six of the seven genes highlighted as differentially expressed between proliferating and resting subpopulations were identified as SV genes (e.g., *THBS1* and *CENPF1*, **Supp Fig. 11C**). This result is also consistent with previous studies which observed that high confluence in cell culture, promoting cell-to-cell communication and crowding, leads to spatial dependency in gene expression^23^. We also considered negative control probes in these data, which were not detected as spatially variable, thereby further confirming the statistical calibration of SpatialDE (**Supp Fig. 11D**).

Herein, we have presented a method for identifying spatially variable genes. The increased availability of high-throughput experiments, including spatially resolved RNA-seq, means that there will be a growing need for methods that account for this new dimension of expression variation, such as SpatialDE.

We applied our model to data from multiple different protocols, from spatial transcriptomics to multiplexed single-molecule FISH, considering both tissue systems and cell lines. The extent of spatial variation we observed in cell lines may be surprising, a result that is consistent with recent studies that have reported coordinated expression changes across neighbouring cells^23^. The method is also applicable to temporal data from time-course experiments to identify genes with dynamic expression trends (**Supp. Fig. 8**), an application of SpatialDE for which previous methods exist^24–26^, but previous methods are likely less computationally efficient than SpatialDE. In principle, SpatialDE can also be applied to 3-dimensional data, e.g., from serial sections of 2-dimensional data, or from *in situ* sequencing when such technologies mature^11,12^.

SpatialDE generalizes previous approaches for the detection of highly variable genes, most notably methods designed for scRNA-seq^13^. Our model separates spatial variation from non-spatial effects, which may include biological and technical variability. Underlying this approach is the assumption that technical noise is independent across sampling positions, which circumvents the need to explicitly model technical sources of variation, which enables applications to virtually any protocol.

Future extension of SpatialDE could be tailored towards specific platforms, for example to make use of spike-in standards, thereby explicitly estimating technical variance components. Other areas of future work are extensions for incorporating information about the tissue makeup or local differences in cell density. Finally, there exist spatial clustering methods that are focused on clustering cell positions rather than genes^15^, which could in the future be combined with the automated expression histology presented here.

## Methods

Methods, including statements of data availability and any associated accession codes and references, are available in the online version of the paper. Full details of the derivation and implementation of SpatialDE are provided in Supplementary Methods.

## Acknowledgements

The authors wish to thank Damien Arnol and Francesco Paolo Casale for helpful advice on statistics and data normalisation. Jeffrey Moffitt helped us understand the data format for available MERFISH data. In addition, we are thankful to Aaron Lun, Martin Hemberg, Daniel Kunz, and Kerstin Meyer for feedback on the manuscript. V.S. was supported by the EMBL International PhD Program, S.A.T. was supported by the Wellcome Trust and ERC Consolidator Grant “ThDEFINE”, O.S. received funding from EMBL core funding, the Wellcome Trust and the EU.

## Author contributions

V.S., O.S., conceived the method. V.S. implemented the method and generated the results. V.S., S.A.T, O.S. interpreted the results. V.S., S.A.T, O.S. wrote the paper.

## Online methods

### SpatialDE model

Spatial DE models the gene expression profiles *y* = (*y*_1_,…, *y*_*N*_) for a given gene across spatial coordinates *X* = (*x*_1_…*x*_*N*_), where each *x*_*n*_ can have arbitrary dimensionality, using a multivariate normal model of the form

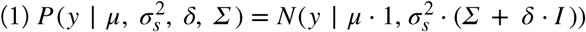

The fixed effect *μ*_*g*_ ⋅ 1 accounts for mean expression level and Σ denotes a spatial covariance matrix, which by default is defined to model general spatial effects using the so called squared exponential covariance function^16^

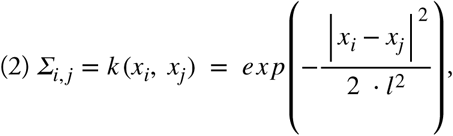

whereby the covariance between pairs of cells *i* and *j* is modelled to decay exponentially with the squared distance between them in the ***X*** coordinates. The hyperparameter *l*, also known as the *characteristic length scale,* determines how rapidly the covariance decays as a function of distance.

The second covariance term *δ* • *I* explains independent non-spatial variation in gene expression, where the ratio FSV = 1 / (1 + *δ*) can be interpreted as the fraction of expression variance attributable to spatial effects. Model parameters are fit using maximum marginal likelihood, an optimisation problem with closed form solutions for the parameters *μ* and α_*s*_, depending on *δ*. Gradient based optimization is used to determine *δ*, and the hyperparameter *l* is determined via grid search. Naïve methods for evaluating the marginal likelihood in Eq. (1) scale cubical in the number of cells, thus prohibiting applications to larger datasets. We adapt algebraic reformulations that have been proposed in statistical genetics^27,28^, coupled with efficient pre-computations of all terms possible, to improve scalability of the model (**Supp. Fig. 1**).

### Statistical significance

To estimate statistical significance, the model likelihood of the fitted SpatialDE model is compared to the likelihood of a model that corresponds to the null hypothesis of no spatial covariance,

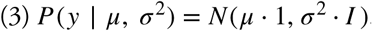

P-values are then estimated analytically based on the *χ*^2^ distribution transformation with one degree of freedom. Multiple testing across genes can be adjusted for using the Q-value method^29^ thereby controlling the false discovery rate (FDR).

### Model selection

Following significance testing, the spatial covariance patterns identified can be further investigated by comparisons of models with alternative covariance functions. In addition to the squared exponential covariance (Eq. (2)), SpatialDE implements comparisons with covariance functions that assumes linear trends as well as periodic signatures (**Figure 1B**), which are compared using the Bayesian information criterion:

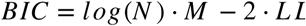

Here *M* denotes the number of hyperparameters of a given model, *N* the number of observations, and *LL* is the log marginal likelihood of the data. For guidance on how to interpret these inferences and alternative functional forms, see **Supp. Methods.**

### Automatic expression histology

To group spatially variable genes with similar spatial expression patterns, SpatialDE implements a clustering model based on the same spatial GP prior as used to test for spatially variable genes (Eq. (1)). Let *Y* = (*y*_1_,…,*y*_*G*_) be the expression matrix of ***G*** spatially variable genes in each spatial location (now each *y*_*g*_ is a vector of *N* observations), ***μ*= {*μ*_1_,.., *μ*_*K*_}** is the matrix of ***K*** underlying patterns, so the vector μ_k_ represents pattern ***k***. Further, let ***Z*** be a binary indicator matrix that assigns genes to patterns. Then the full model can be written as:

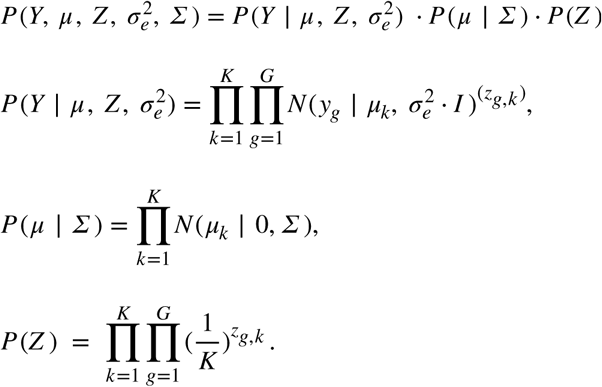

The parameter 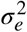 is the noise level for the model, and ***Σ*** is the spatial covariance matrix defined based on spatial coordinates (see Eq. (2)). This model can be regarded as an extension of the classical Gaussian mixture model^30^, with the addition of a spatial prior on cluster centroids. The posteriors of *μ* and *Z* are approximated using variational inference^30^, while the noise level 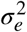 is estimated by maximising the variational lower bound. The length scale *l* for the covariance ***Σ*** is specified by the user, as is the number of fitted patterns, ***K***. The choice of *l* can be informed by the fitted length scales in the SpatialDE significance test. See **Supp. Methods** for details on inference and derivation of variational updates.

After inference, the posterior expectations *<u>μ</u>* and *<u>Z</u>* of the parameters can be used to visualise any histological pattern through plotting <u>*μ*_*k*_</u> over the *x* coordinates. The most likely assignment of genes to an individual pattern is determined by the largest value in the vector <u>*Z*_*g*_</u>, which corresponds to the posterior probabilities of a gene belonging to each pattern.

### Relationship to prior work

SpatialDE is related to a number of existing methods based on Gaussian processes. First used in geostatistics^31^, GP models have been applied to test for differential gene expression over time^32^, including the analysis of bifurcation points^26^, and general tests for temporal variability^25,33–35^.

We have here adapted GP models to spatial transcriptome data, although the model can also be applied to univariate data (**Supp. Fig. 12**) or higher-dimensional inputs. The main technical innovations presented here are three-fold. First, the model presented is faster than existing methods, by leveraging computational tricks previously proposed in the context of statistical genetics (**Supp. Fig. 1**, Section above). Second, we combine spatial GPs with model selection using BIC, a criterion that has been used for time series analyses^36^ but has not previously been considered in this context. Third, we propose an efficient and versatile spatial clustering model based on the same statistical framework.

### Availability of code and data

SpatialDE is implemented in Python 3.5. An open source implementation is available fromhttps://github.com/Teichlab/SpatialDE together with a Stan version, and can be installed from PyPI using the command ‘pip install spatialde’. The release includes tutorials and example vignettes for reproducing the presented analyses, as well as all pre-processed datasets considered in this study. An R-based Bioconductor implementation is in preparation.

### Data sets and processing

#### Spatial Transcriptomics data

The count tables from Stahl et al^7^ were downloaded from the website http://www.spatialtranscriptomicsresearch.org/datasets/doi-10-1126science-aaf2403, linked from the publication. For the breast cancer data, we used the file annotated as “Layer 2” with the corresponding HE image. For the mouse olfactory bulb, we used the file named “Replicate 11” with corresponding HE image. Images included in figures were cropped, down-scaled and converted to grayscale to conserve file sizes. When performing automatic expression histology, the number of patterns was set to 5 for both data sets, the characteristic length scale was set to 105 μm for the breast cancer data, and to 150 μm for the olfactory bulb data.

#### SeqFISH data

We downloaded the expression table from the supplementary material of Shah *et al*^8^ and extracted cell counts from the region annotated with number 43 in the 249-gene experiment (Table S8 in the original publication). The shape of the data suggested this corresponded to a region in the lower left part of the corresponding supplementary figure, informing the schematic shown in **Figure. 2D** (only used for the purpose of illustration). In the automatic histology analysis, the number of patterns was set to 5, and the characteristic length scale was set to 50 μm.

#### MERFISH data

From the website http://zhuang.harvard.edu/merfish we downloaded the file “data for release.zip” which contain data from Moffitt et al^9^ We used the files in the folder called “Replicate 6”, as these had the largest number of cells and highest confluency.

#### Frog development RNA-seq data

We downloaded the TPM expression table for Clutch A from GEO accession GSE65785 which was referenced in the original publication^24^.

### Expression count normalisation

The SpatialDE model is based on the assumption of normally distributed residual noise and independent observations across cells. To meet these requirements with spatial expression count data we have identified two normalisation steps (**Supp. Methods**). First, we use a variance stabilising transformation for negative binomial distributed data to satisfy the first condition known as Anscombe’s transformation. Second, we noticed that generally the expression level of a given gene correlates with the total count in a cell / spatial location. To ensure that SpatialDE captures the spatial covariance for each gene beyond this effect, log total count values are regressed out from the Anscombe-transformed expression values before fitting the spatial models.

## Supplementary information

### Supp. Methods

Full derivation of the SpatialDE model, detailed descriptions significance test, model selection, dataset processing, expression normalisation. As well as full details on the automatic expression histology model.

### Supp. Table 1

Supplementary table with SpatialDE analysis results; Each tab contains the output of an analysis. Tabs start with a name of the dataset: “BC” for breast cancer data; “Frog” for frog development time course; “MERFISH” for MERFISH cell culture data; “MOB” for mouse olfactory bulb data; and “SF” for SeqFISH hippocampus data. Tabs named “NAME]_final_results” indicates SpatialDE significance test results for each gene. Tabs named “[NAME]_MS_results” refer to model selection results for significant genes. The tabs named “[NAME]_AEH_results” indicate the assignment of genes to inferred histological patterns using the AEH analysis. The functional enrichment analysis results for the breast cancer data are in the tab “BC_gProfiler”.

**Supp. Fig 1.**
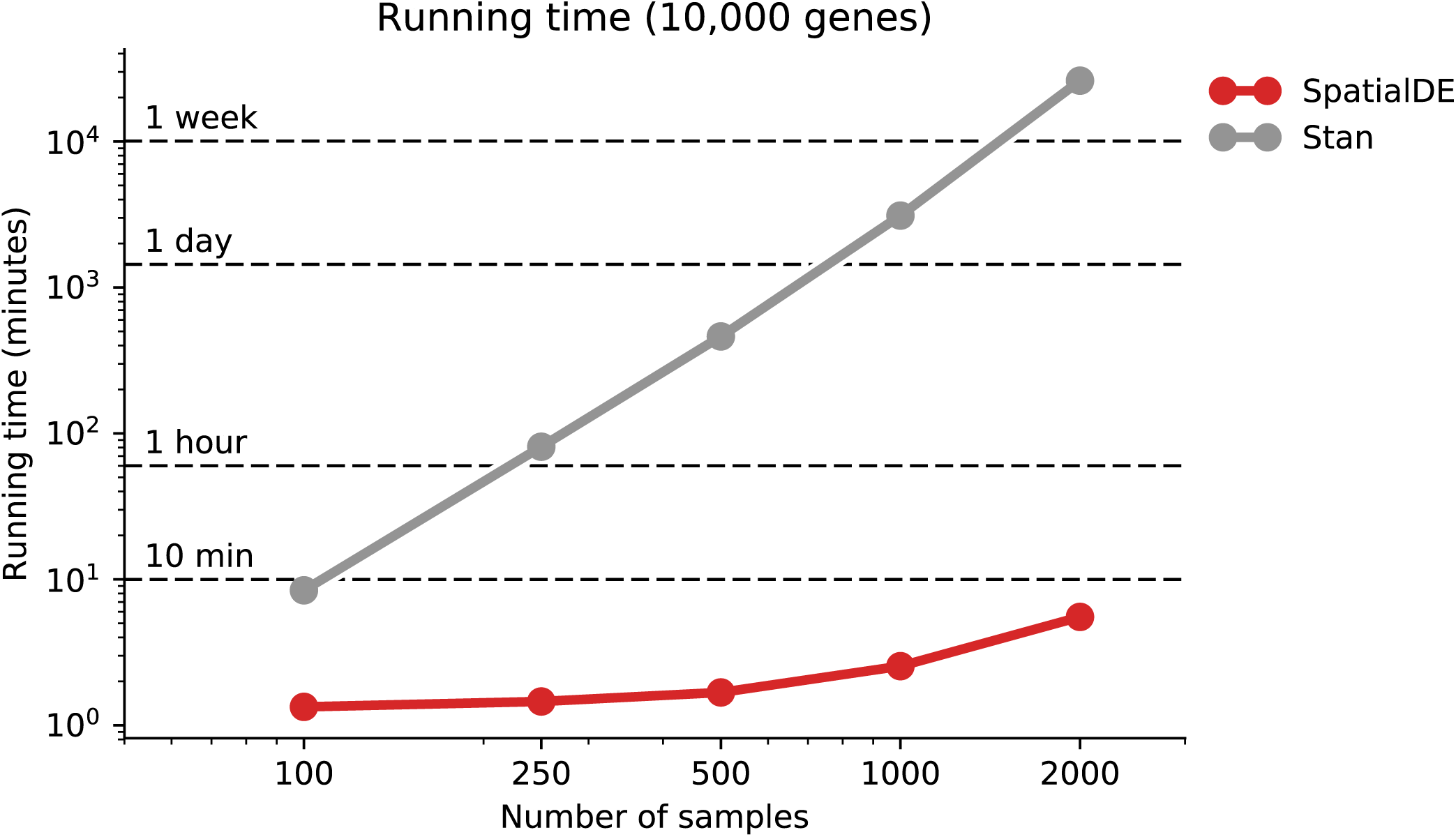
Computational efficiency of SpatialDE. Compared is the SpatialDE native implementation versus a Stan implementation of the same model. Caching operations and linear algebra speedups are used where possible, enabling tractable genome-wide analyses with thousands of samples. Benchmarks performed on a late 2013 iMac with 3.2 GHz Intel Core i5 processor.

**Supp. Fig. 2.**
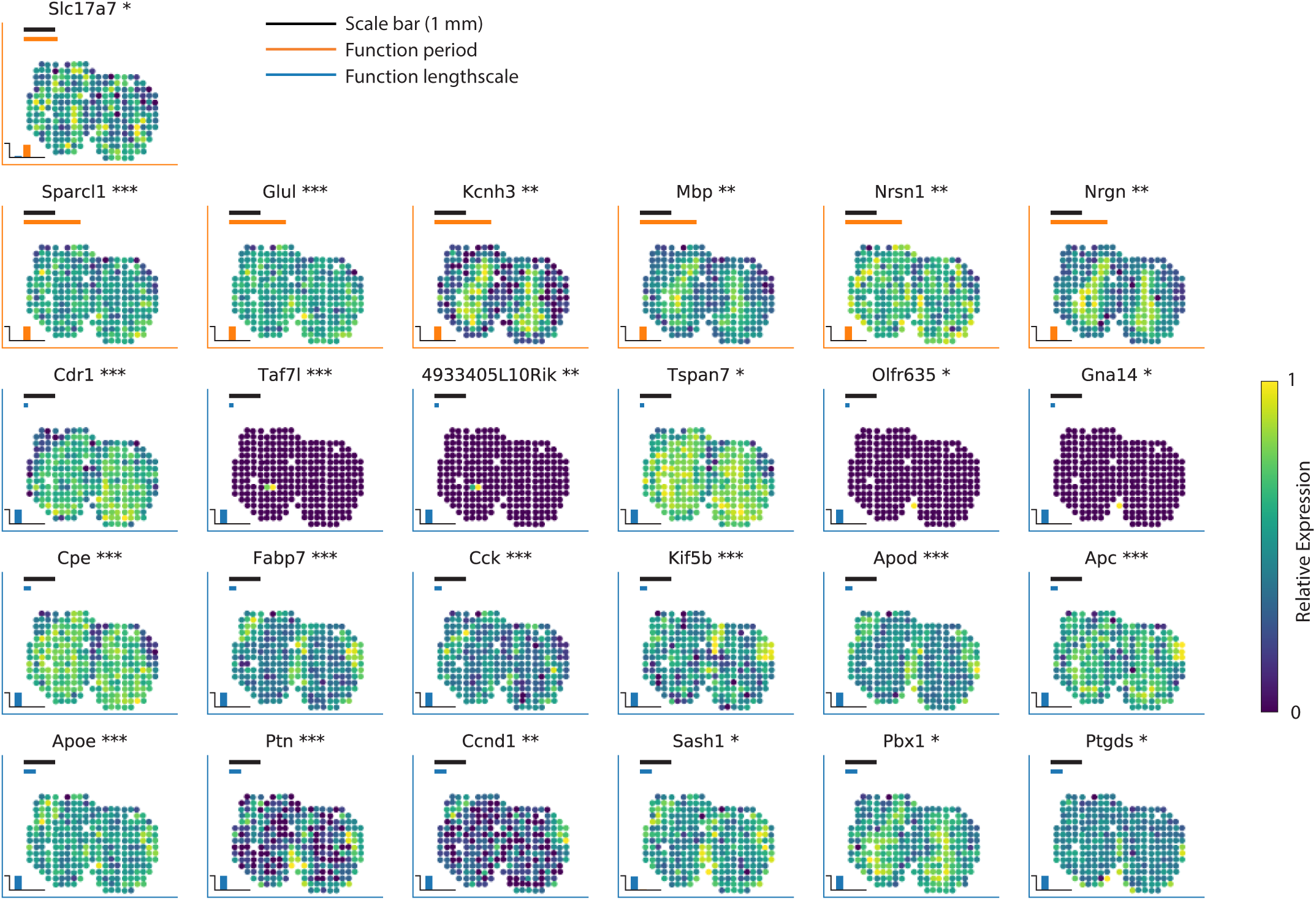
Expanded example of spatially variable genes identified in the mouse olfactory bulb. Additional spatial expression patterns for 25 SV genes (out of 67, FDR<0.05), selected to represent expression patterns with different function periods and length scales.

**Supp. Fig. 3.**
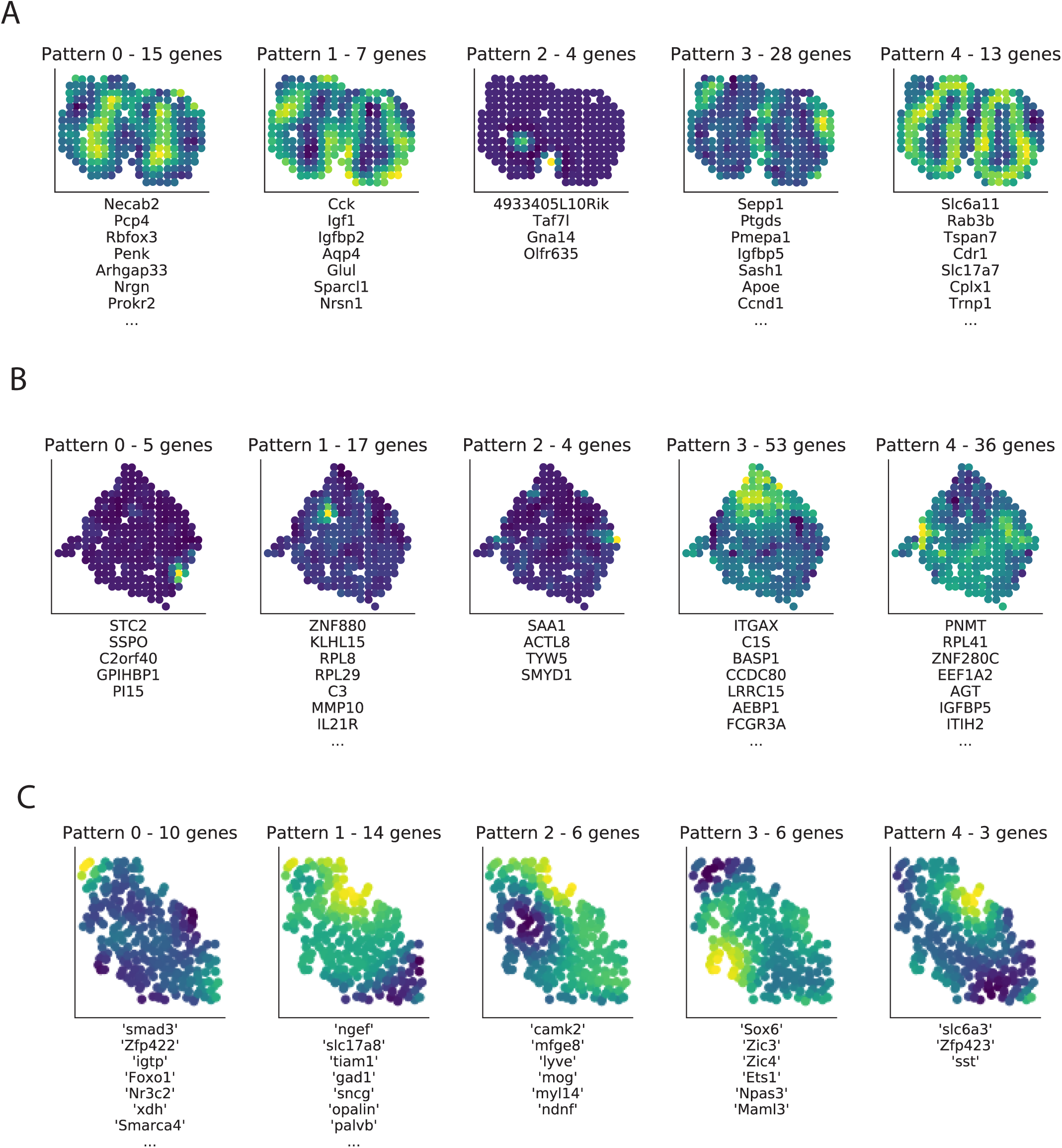
Results from automatic expression histology. **(A)** Applied to spatial transcriptomics data of mouse olfactory bulb, for K=5 spatial expression patterns. Color denotes the expression level of the inferred spatial pattern. The number of genes assigned to each pattern is written in the title of each plot, with representative gene assignments listed below each panel. Genes linked to the same pattern can be thought of as spatially coexpressed. **(B)** Same as **A** but applied to spatial transcriptomics data from breast cancer biopsy. **(C)** As in **A** but for SeqFISH data from mouse hippocampus.

**Supp. Fig. 4.**
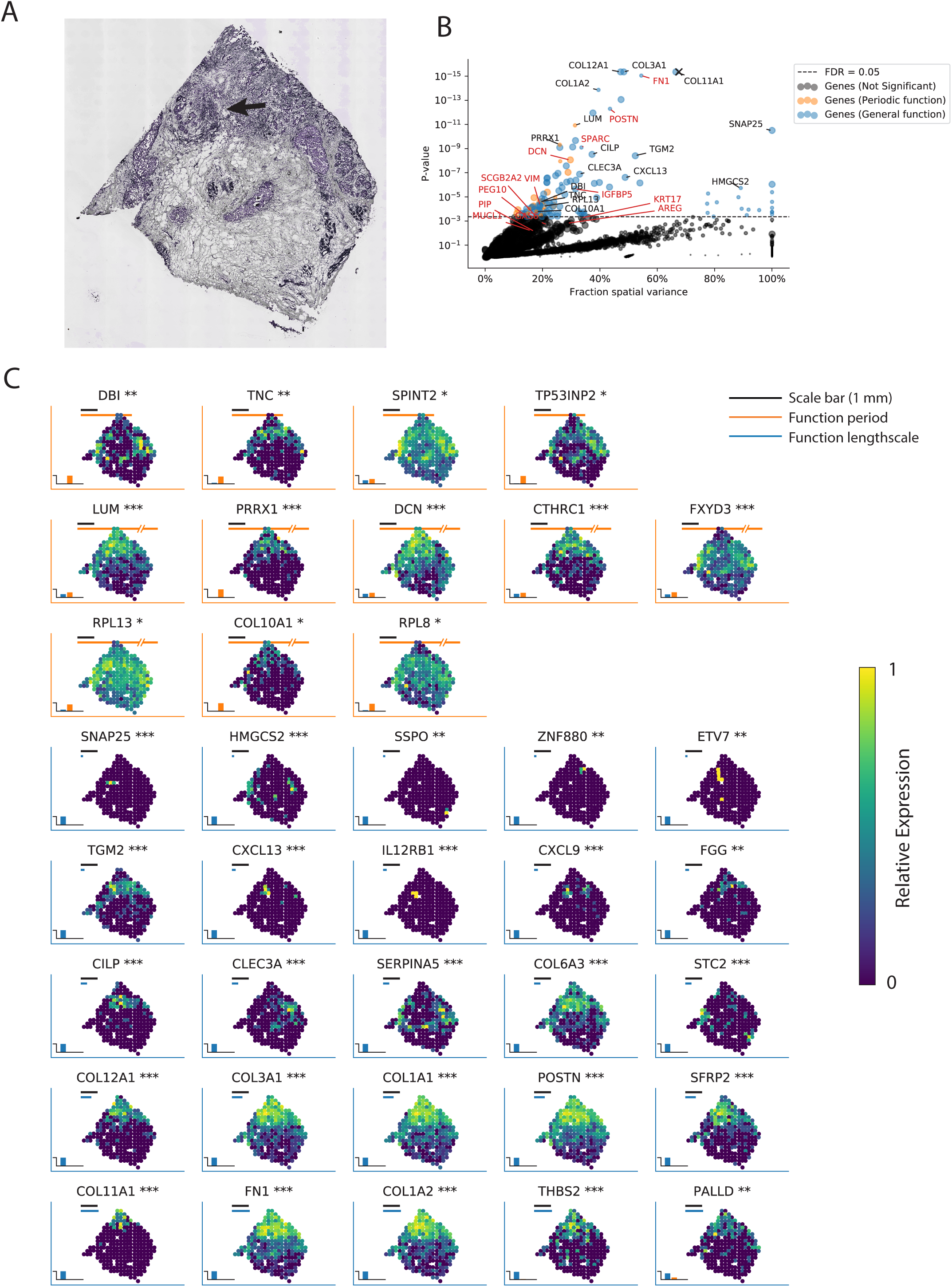
SpatialDE applied to breast cancer biopsy tissue. (**A**) Corresponding HE image of breast cancer tissue from spatial transcriptomics. (**B**) Fraction of variance explained by spatial variation (X-axis) versus P-value (y-axis) for all genes. Dashed line corresponds to the FDR=0.05 significance threshold (N=115 genes). Genes classified as periodically variable are shown in orange (N=22), genes with a general spatial dependency in blue (N=93). Disease-implicated genes annotated based on prior knowledge (Stahl et al.^7^) are highlighted with red labels. Other representative genes of various function periods or length scales are annotated with black labels. The X symbol shows the result of applying SpatialDE to the estimated total RNA content per spot. (**C**) Visualization of 37 selected spatially variable genes with different periods and length scales. The black scale bar corresponds to 1 mm. Colors and labels as in (Figure 2D).

**Supp. Fig. 5.**
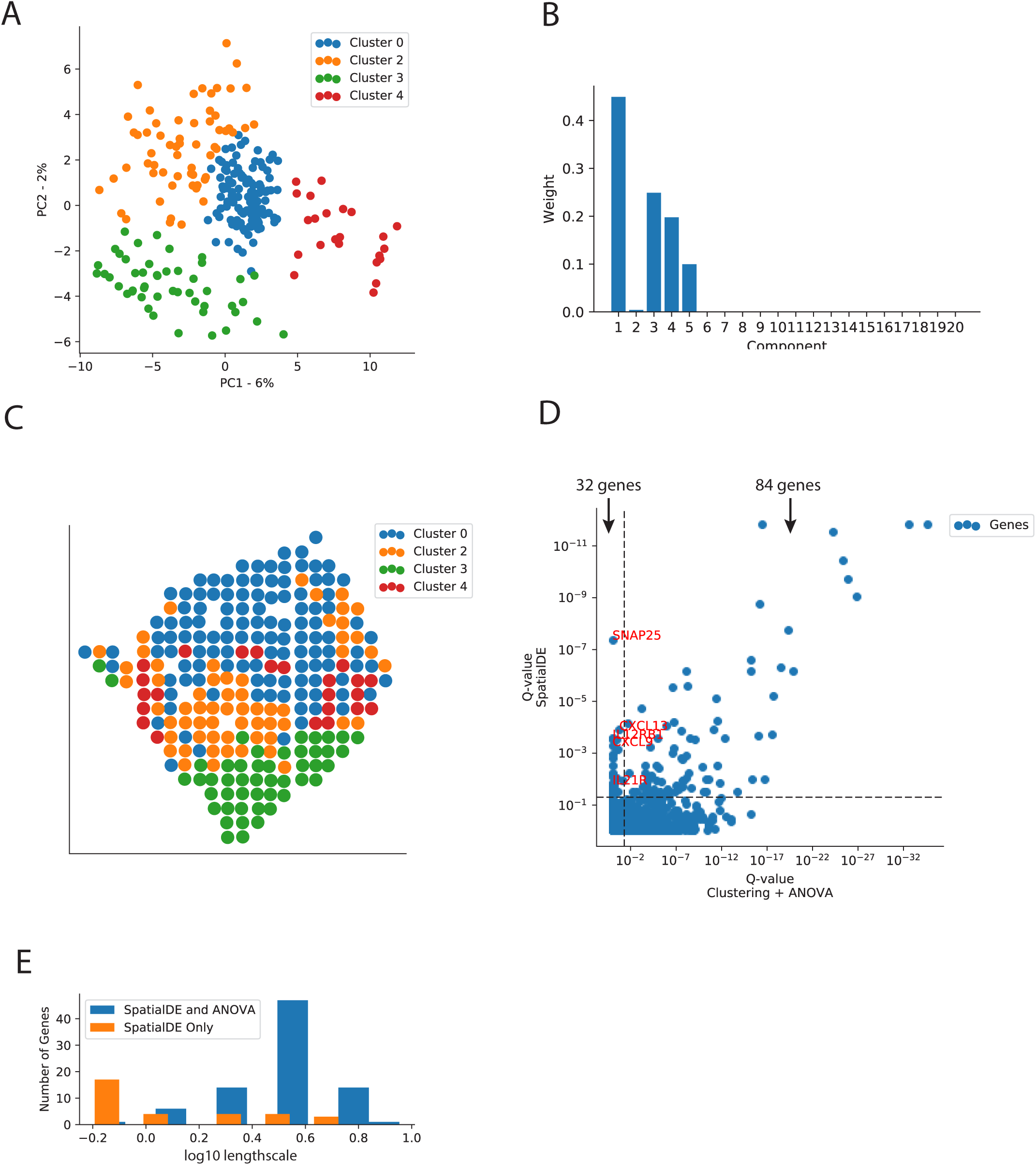
Comparison to differential expression analysis using unsupervised clustering of spots. **(A)** Principal component analysis of individual “spots” from the spatial transcriptomics breast cancer data, color coded by cluster membership for N=4 clusters (identified by Bayesian Gaussian Mixture Modelling). **(B)** Bayesian Gaussian Mixture Model cluster probabilities, discretizing the 250 spatial breast cancer “spots” into four clusters (analysis ignores spatial structure). **(C)** Visualization of cluster membership in the original tissue context. **(D)** Scatter plot of P-values from an ANOVA test between clusters (x-axis) versus significance of spatial variation using SpatialDE (y-axis). 83 genes were identified as significantly variable by both approaches (FDR<0.05); 32 genes are significant only in the SpatialDE test, among them immune genes. **(E)** Histogram of the fitted length scales for SV genes detected by both approaches (blue) and SV genes exclusively detected by SpatialDE (orange). Genes that were only detected by SpatialDE were associated with smaller length scales, indicating localized expression patterns.

**Supp. Fig. 6.**
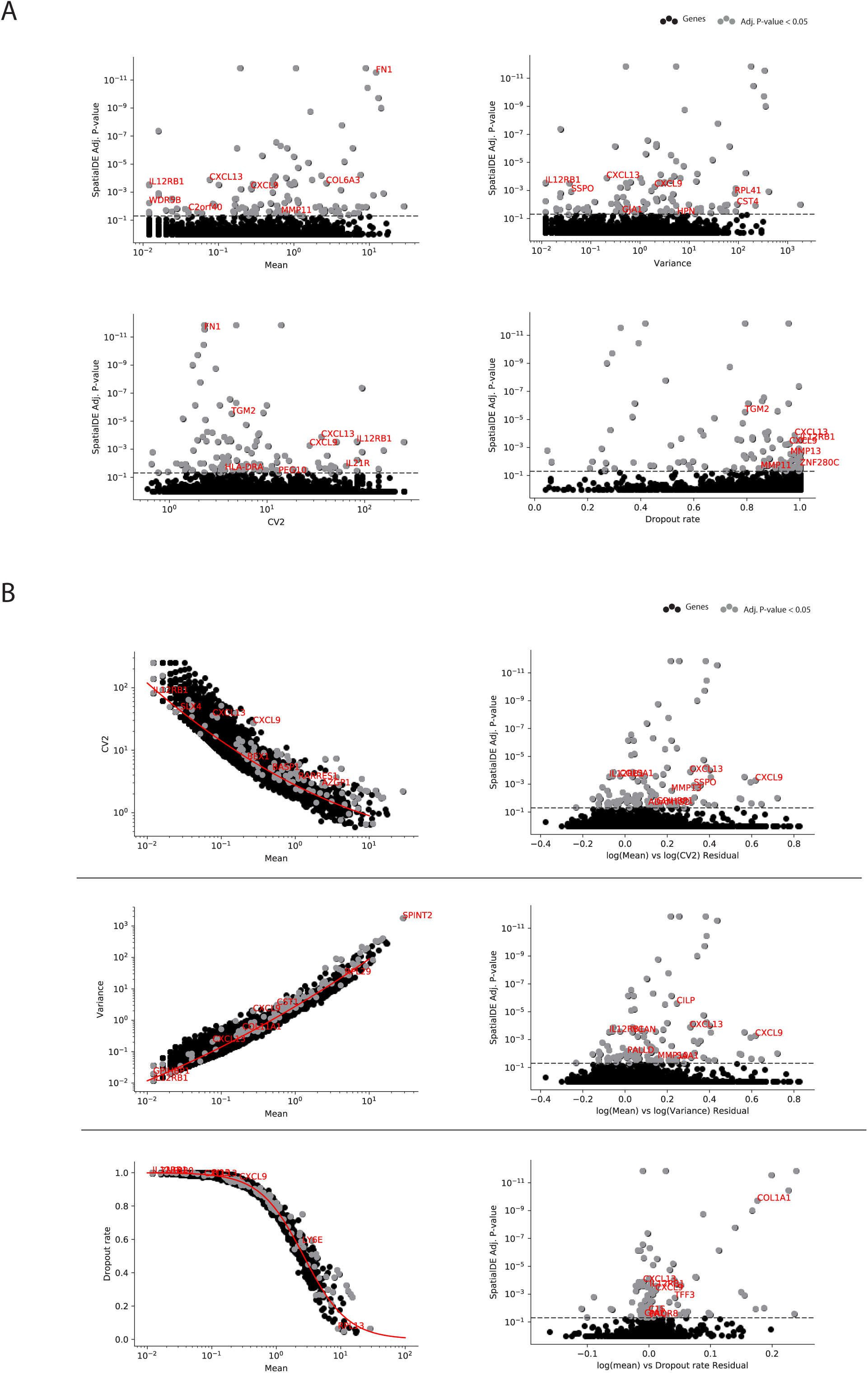
Comparison of SpatialDE to alternative measures of expression heterogeneity in the breast cancer tissue. **(A)** Comparison of the SpatialDE test (P-values, y-axis) versus commonly used statistics (x-axis) - Upper left: Mean, Upper right: Variance, Lower left: CV2 (squared coefficient of variation), Lower right: Dropout rate (fraction of cells/samples a gene is not detected in). Random selection of significant SV genes highlighted in red for context. No dependence between SpatialDE significance levels and expression level (mean) or variance was observed. **(B)** Comparison of significance of SV for genes identified by SpatialDE versus commonly used strategies for defining highly variable genes based on regression models between summary statistics: Relation with CV2 (Upper) or Variance (Middle), or with dropout fraction (Bottom). The rightmost column of plots shows residuals compared with the SpatialDE significance; polynomial regression for CV2 and Variance, logistic regression for dropout rate. Significant SV genes as identified by SpatialDE (FDR<0.05) are shown in grey. Other, non-significant genes are shown in solid black. The SV genes significance are orthogonal to HVG measures, indicating that spatial variation is different from general variability.

**Supp. Fig. 7.**
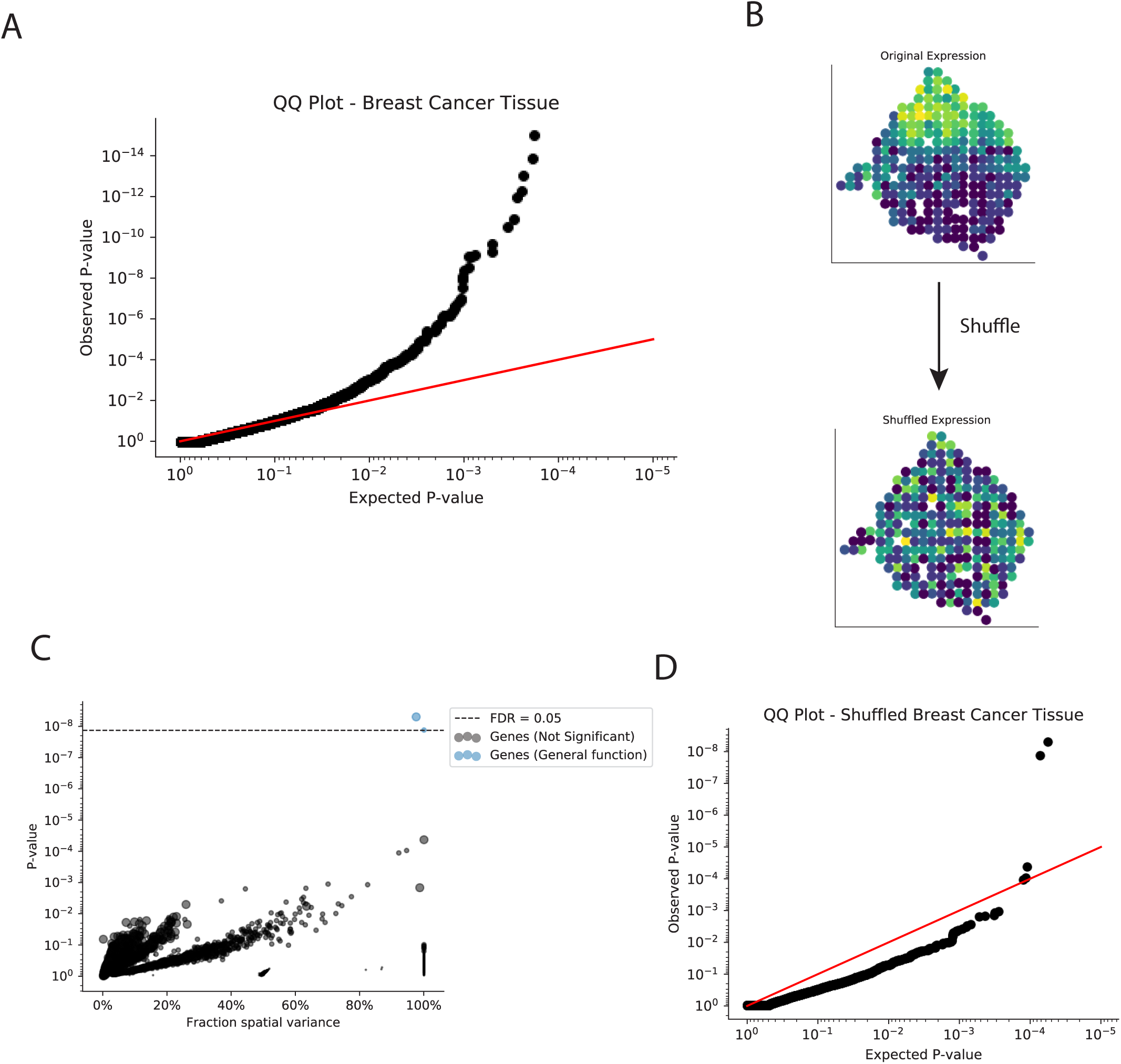
Assessment of statistical calibration of SpatialDE through data randomization. **(A)** QQ-plot of expected P-values (Chi2 distribution with 1 degree of freedom) versus observed P-values from the SpatialDE tests on the breast cancer data. **(B)** To simulate data from an empirical null, without spatial structure, expression values were shuffled among the sampled coordinates. Shown is the resulting expression pattern for *COL3A1* expression, as representative example. **(C)** P-values for genes on shuffled data, with the number of detected SV genes (FD<0.05) being consistent with the selected false discovery rate. **(D)** Analogous QQ-plot as in **A** on shuffled expression values. P-values follow the null distribution, indicating that the model is calibrated.

**Supp. Fig. 8.**
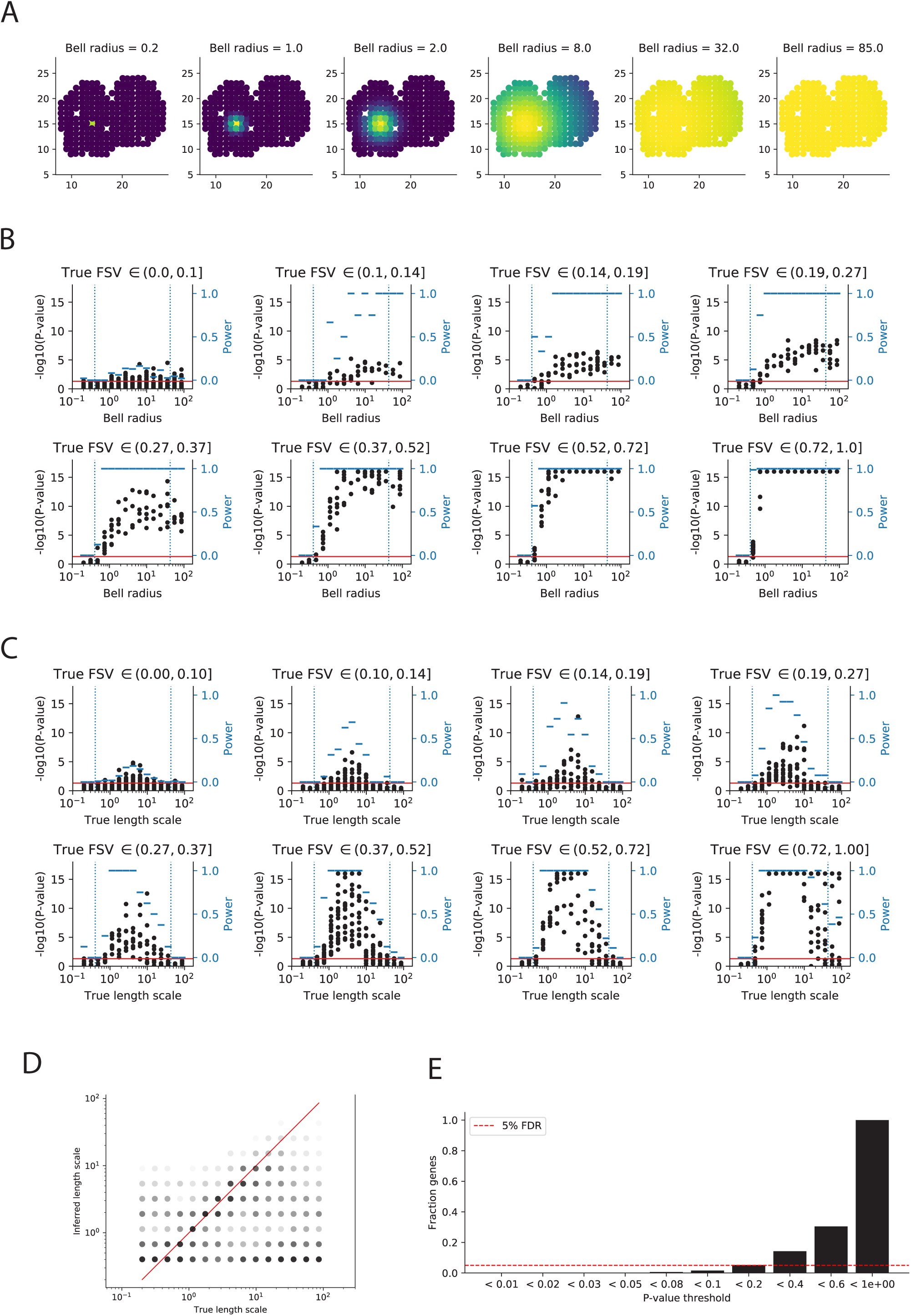
Assessing statistical calibration of SpatialDE through simulations. **(A-B)** Simulation of bell curve shaped data on the mouse olfactory bulb coordinates. **(A)** Example of six bell curves with different radii. **(B)** Results from applying SpatialDE to data from 3,000 bell curves with different radii and stratified over different levels of simulated noise (fraction of spatial variance, FSV). Red line denotes P=0.05 significance level, vertical blue dotted lines indicate smallest and largest pairwise distances observed in mouse olfactory bulb data. Black dots denote -log10(P-values) (left axis), while blue dashes indicating statistical power for detecting true simulated SV genes (fraction true positives) for each bell radius (right axis). **(C)** Analogous results as in **B**, however when simulating data from the generative model underlying SpatialDE, considering different values of FSV and length scales (3,000 simulations). **(D)** Scatter plot of inferred length scales (y-axis) versus simulated length scale (x-axis) for the data shown in **C**. **(E)** Simulation of 3,000 genes from the null model with no spatial covariance. Bars denote the fraction of significant SV genes at different P-value thresholds (x-axis). The proportion of false positive genes was lower than the controlled FDR, indicating that the test is conservative.

**Supp. Fig. 9.**
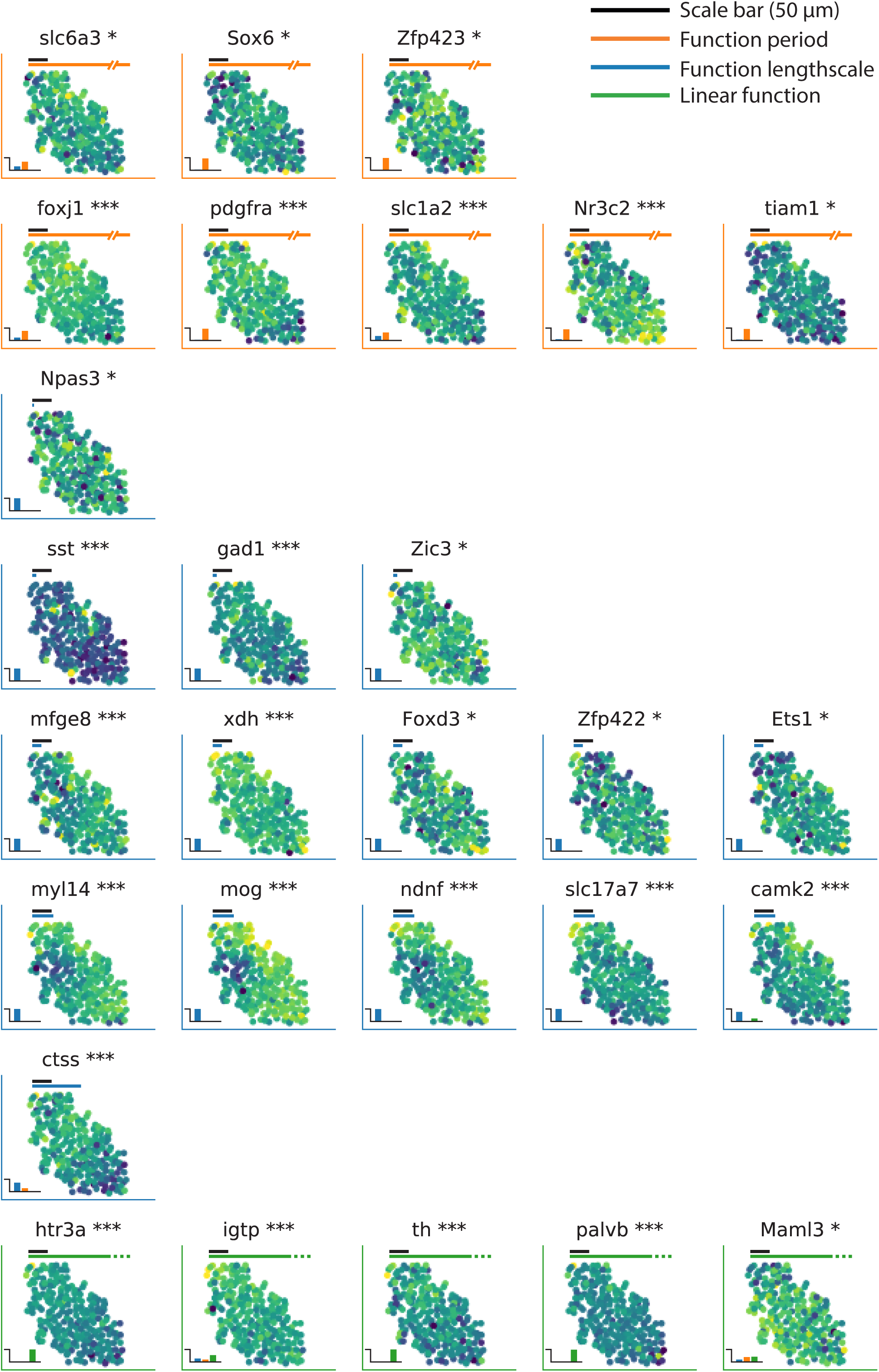
Expanded examples of spatially variable genes for the mouse hippocampus dataset. Visualization of 28 SV genes (out of 249, FDR<0.05) from the mouse hippocampus SeqFISH data, showing selected genes with periodic, linear, and general spatial dependencies with different estimated length scales. Black scale bar corresponds to 50 μm. Colors and labels as in (Figure 2B).

**Supp. Fig. 10.**
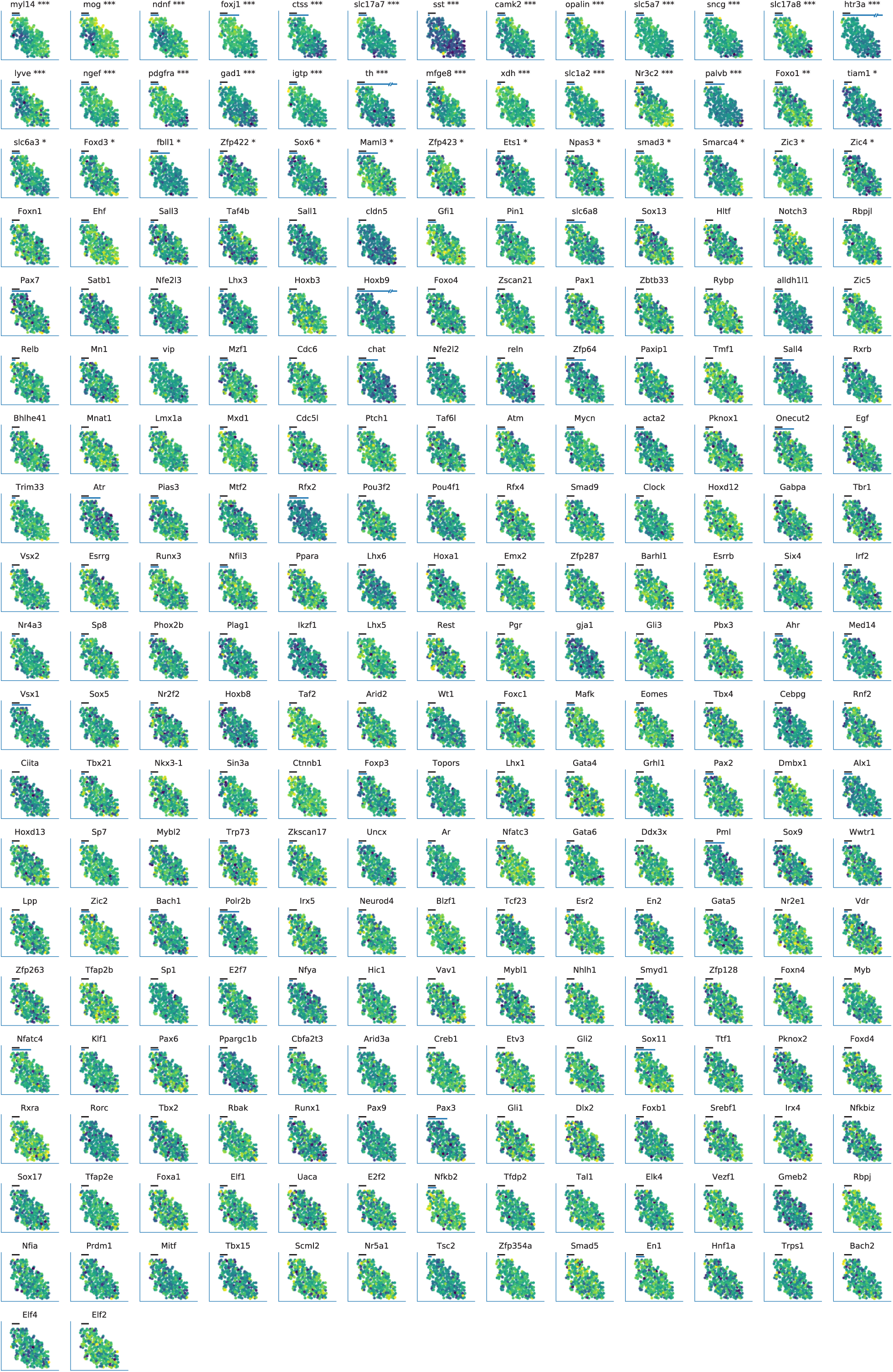
Visual inspection of genes from SeqFISH data. All 249 genes measured in the SeqFISH data. Plots are ordered left to right then top to down with increasing P-values. Stars next to gene names denote significance levels (* FDR < 0.05, ** FDR < 0.01, *** FDR < 0.001).

**Supp. Fig. 11.**
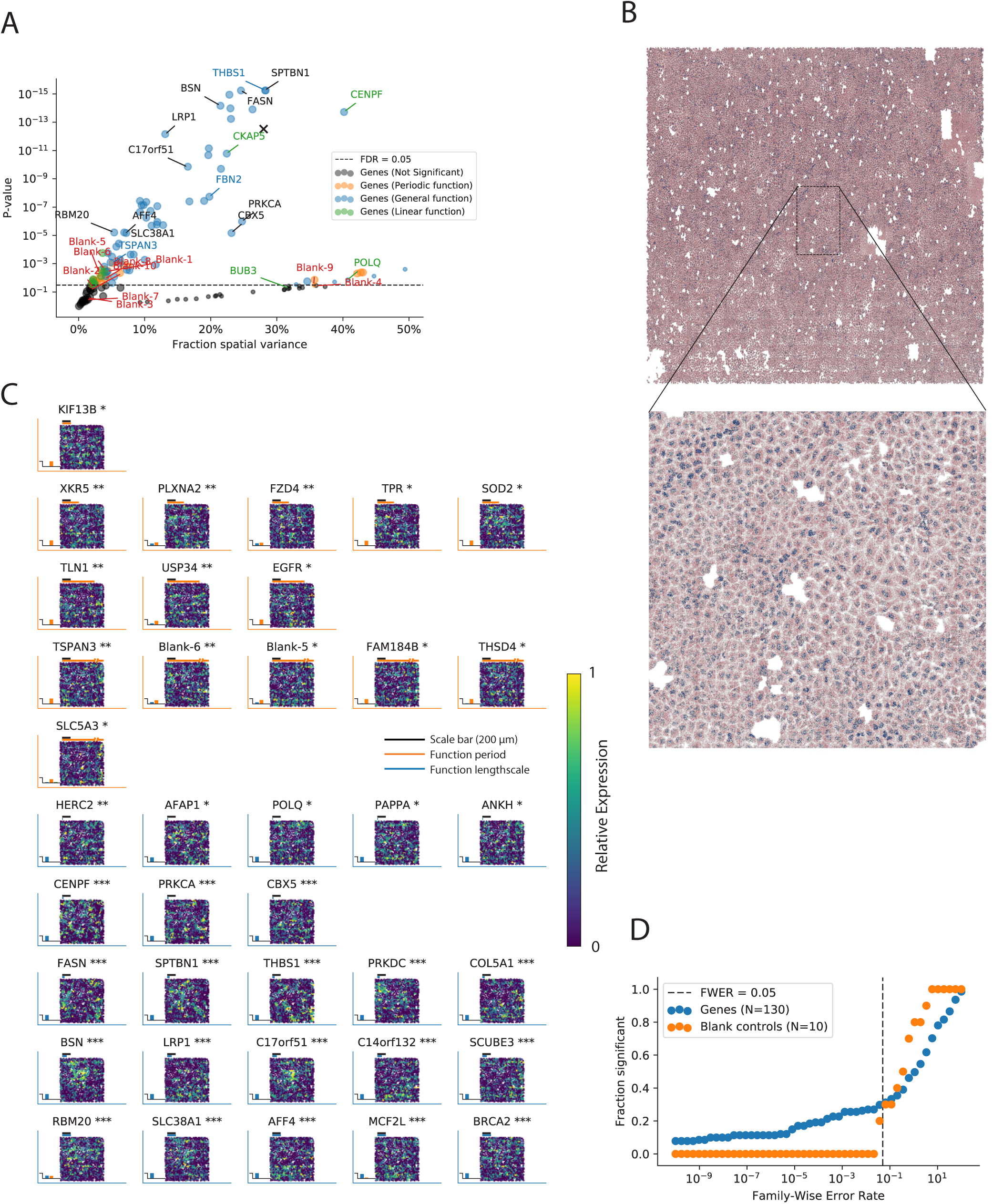
Application to MERFISH data. **(A)** In a MERFISH study of an osteosarcoma cell culture of 139 probes from Moffitt *et al,* SpatialDE identified the majority of the probes as spatially variable (66%, FDR<0.05). 21 of 92 significant SV genes were assigned to a periodic function by the model, and 9 genes had linear functions. Red labels indicate negative control probes. Genes indicated as enriched in proliferating cells in the original study are marked in green, and depleted genes in blue. **(B)** Visualization of the MERFISH data by plotting general RNA probes in pink and MALAT1 probes in blue on two 512 x 512 virtual pixel grids at different scales. The original imaged region was 5.2 mm wide and 8.2 mm high totalling 38,594 cells (upper). We analysed a region of 1 mm x 1 mm in the middle of the cell culture with 1,056 cells (lower). **(C)** Expression levels in the cell culture region visualised for selected SV genes with various fitted periods and length scales (Significance levels and colors as in Figure 2). Black scale bar corresponds to 200 μm. **(D)** Fraction of gene probes and control probes detected as significant SV genes as a function of the family-wise error rate (FWER). The number of significant control probes was in line with the FWER.

**Supp. Fig. 12.**
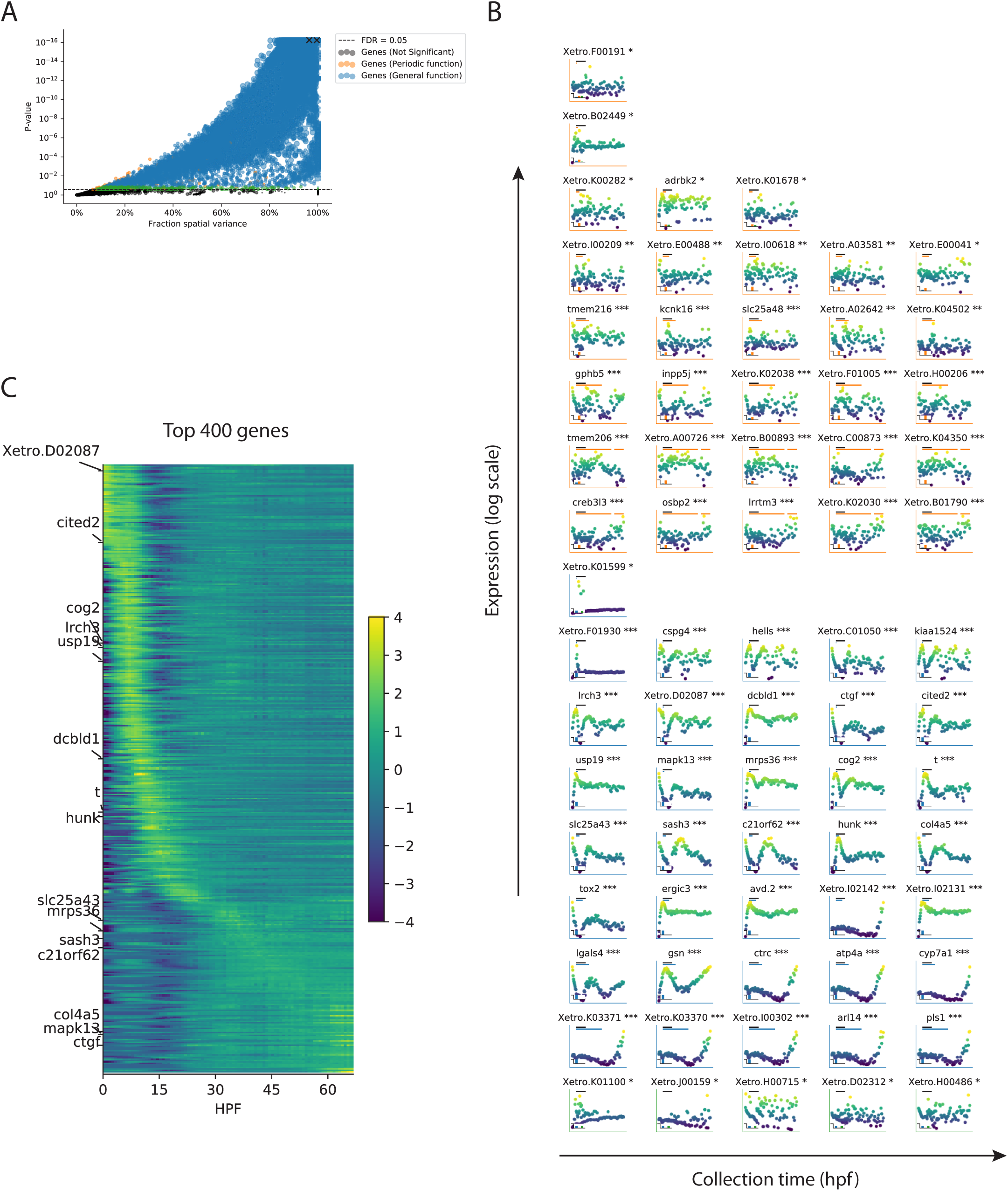
Application to a gene expression time-course data set. **(A)** SpatialDE applied to a developmental time course (89 time points from Owens *et al),* identifying the majority of genes as spatially variable (21,009 out of 22,256 genes, FDR < 0.05). Of these, 241 were assigned to periodic patterns, and 269 were detected with linear trends. Colors and point sizes as in Figure 2. The X marks indicates result of running SpatialDE on RNA spike-in content and the number of detected genes, proxies for the RNA content in the embryos. **(B)** Examples of temporally variable genes of various periods and length scales. Black scale bar corresponds to 12 hours in the time-course, periods and length scales of functions are indicated relative to this. Collection time in units of hours post fertilization (hpf) **(C)** The expression patterns of the top 400 significantly SV genes are visualised, ordered by the time they reach their highest expression value. Example genes from **B** are annotated.

**Supp. Fig. 13.**
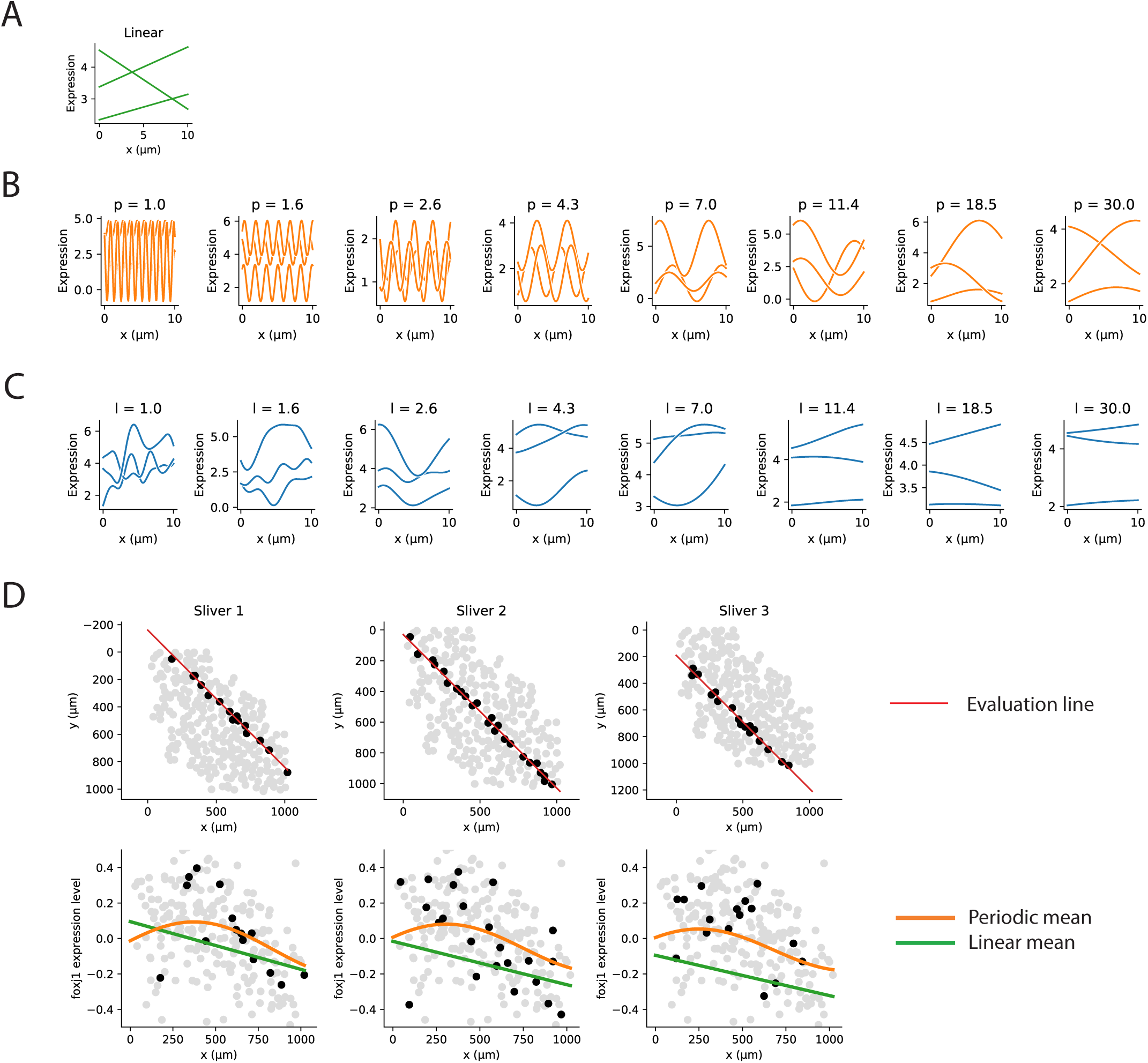
Interpretation of periods and length scales. **(A-C)** Simulated noise free 1D data with different covariance functions. **(A)** Three simulations with linear covariance. **(B)** Simulations from periodic covariance functions with different period values. **(C)** Simulations from squared exponential covariance function with different characteristic length scales. **(D)** Visualization of a gene with very long inferred period length. Top row shows the region where the Gaussian processes are predicted. Bottom row shows expression level on the y-axis instead of the spatial y-coordinate. Curves show expression level prediction from the Gaussian process along the red line indicated in the top panel. Cells in grey or black are spatially closer to the red line.

## References

1. Ledford, H. The race to map the human body - one cell at a time. Nature 542, 404–405 (2017).

2. Lee, J. H. Quantitative approaches for investigating the spatial context of gene expression. Wiley Interdiscip. Rev. Syst. Biol. Med. 9, (2017).

3. Achim, K. et al. High-throughput spatial mapping of single-cell RNA-seq data to tissue of origin. Nat. Biotechnol. 33, 503–509 (2015).

4. Satija, R., Farrell, J. A., Gennert, D., Schier, A. F. & Regev, A. Spatial reconstruction of single-cell gene expression data. Nat. Biotechnol. 33, 495–502 (2015).

5. Junker, J. P. et al. Genome-wide RNA Tomography in the zebrafish embryo. Cell 159, 662–675 (2014).

6. Chen, J. et al. Spatial transcriptomic analysis of cryosectioned tissue samples with Geo-seq. Nat. Protoc. 12, 566–580 (2017).

7. Ståhl, P. L. et al. Visualization and analysis of gene expression in tissue sections by spatial transcriptomics. Science 353, 78–82 (2016).

8. Shah, S., Lubeck, E., Zhou, W. & Cai, L. In Situ Transcription Profiling of Single Cells Reveals Spatial Organization of Cells in the Mouse Hippocampus. Neuron 92, 342–357 (2016).

9. Moffitt, J. R. et al. High-throughput single-cell gene-expression profiling with multiplexed error-robust fluorescence in situ hybridization. Proc. Natl. Acad. Sci. U. S. A. 113, 11046–11051 (2016).

10. Ke, R. et al. In situ sequencing for RNA analysis in preserved tissue and cells. Nat. Methods 10, 857–860 (2013).

11. Lee, J. H. et al. Highly multiplexed subcellular RNA sequencing in situ. Science 343, 1360–1363 (2014).

12. Lee, J. H. et al. Fluorescent in situ sequencing (FISSEQ) of RNA for gene expression profiling in intact cells and tissues. Nat. Protoc. 10, 442–458 (2015).

13. Brennecke, P. et al. Accounting for technical noise in single-cell RNA-seq experiments. Nat. Methods 10, 1093–1095 (2013).

14. Vallejos, C. A., Marioni, J. C. & Richardson, S. BASiCS: Bayesian Analysis of Single-Cell Sequencing Data. PLoS Comput. Biol. 11, e1004333 (2015).

15. Pettit, J.-B. et al. Identifying cell types from spatially referenced single-cell expression datasets. PLoS Comput. Biol. 10, e1003824 (2014).

16. Rasmussen, C. E. & Williams, C. Gaussian Processes for Machine Learning, Model Selection and Adaptation of Hyperparameters, Chapter 5. (2006).

17. Lippert, C. et al. FaST linear mixed models for genome-wide association studies. Nat. Methods 8, 833–835 (2011).

18. Jahn, R., Takamori, S., Rhee, J. S. & Rosenmund, C. 10.1038/35025070. Nature 407, 189–194 (2000).

19. Seewaldt, V. L. Cancer: Destiny from density. Nature 490, 490–491 (2012).

20. Reimand, J. et al. g:Profiler-a web server for functional interpretation of gene lists (2016 update). Nucleic Acids Res. 44, W83–9 (2016).

21. Andrews, T. S. & Hemberg, M. Modelling dropouts allows for unbiased identification of marker genes in scRNASeq experiments. bioRxiv (2016).

22. Chen, K. H., Boettiger, A. N., Moffitt, J. R., Wang, S. & Zhuang, X. RNA imaging. Spatially resolved, highly multiplexed RNA profiling in single cells. Science 348, aaa6090 (2015).

23. Battich, N., Stoeger, T. & Pelkmans, L. Control of Transcript Variability in Single Mammalian Cells. Cell 163, 1596–1610 (2015).

24. Owens, N. D. L. et al. Measuring Absolute RNA Copy Numbers at High Temporal Resolution Reveals Transcriptome Kinetics in Development. Cell Rep. 14, 632–647 (2016).

25. Macaulay, I. C. et al. Single-Cell RNA-Sequencing Reveals a Continuous Spectrum of Differentiation in Hematopoietic Cells. Cell Rep. 14, 966–977 (2016).

26. Lönnberg, T. et al. Single-cell RNA-seq and computational analysis using temporal mixture modeling resolves TH1/TFH fate bifurcation in malaria. Science Immunology 2, eaal2192 (2017).

27. Lippert, C. et al. FaST linear mixed models for genome-wide association studies. Nat. Methods 8, 833–835 (2011).

28. Zhou, X. & Stephens, M. Genome-wide efficient mixed-model analysis for association studies. Nat. Genet. 44, 821–824 (2012).

29. Storey, J. D. & Tibshirani, R. Statistical significance for genomewide studies. Proc. Natl. Acad. Sci. U. S. A. 100, 9440–9445 (2003).

30. Bishop, C. M. Pattern recognition and machine learning. (springer, 2006).

31. Krige, D. G. A statistical approach to some basic mine valuation problems on the Witwatersrand. J. South Afr. Inst. Min. Metall. 52, 119–139 (1951).

32. Stegle, O. et al. A robust Bayesian two-sample test for detecting intervals of differential gene expression in microarray time series. J. Comput. Biol. 17, 355–367 (2010).

33. Kalaitzis, A. A. & Lawrence, N. D. A simple approach to ranking differentially expressed gene expression time courses through Gaussian process regression. BMC Bioinformatics 12, 180 (2011).

34. Äijö, T. et al. Methods for time series analysis of RNA-seq data with application to human Th17 cell differentiation. Bioinformatics 30, i113–20 (2014).

35. Eckersley-Maslin, M. A. et al. MERVL/Zscan4 Network Activation Results in Transient Genome-wide DNA Demethylation of mESCs. Cell Rep. 17, 179–192 (2016).

36. Lloyd, J. R., Duvenaud, D., Grosse, R., Tenenbaum, J. B. & Ghahramani, Z. Automatic Construction and Natural-Language Description of Nonparametric Regression Models. arXiv [stat.ML] (2014).

